# Mitochondrial pathway polygenic risk scores are associated with Alzheimer’s Disease

**DOI:** 10.1101/2020.11.06.371286

**Authors:** Devashi Paliwal, Tim W. McInerney, Judy Pa, Russell H. Swerdlow, Simon Easteal, Shea J. Andrews, for the Alzheimer’s Disease Neuroimaging Initiative

## Abstract

**INTRODUCTION:** Genetic, animal and epidemiological studies involving biomolecular and clinical endophenotypes implicate mitochondrial dysfunction in Alzheimer’s disease (AD) pathogenesis. Polygenic risk scores (PRS) provide a novel approach to assess biological pathway-associated disease risk by combining the effects of variation at multiple, functionally related genes.

**METHODS:** We investigated associations of PRS for genes involved in 12 mitochondrial pathways (pathway-PRS) related to AD in 854 participants from Alzheimer’s Disease Neuroimaging Initiative.

**RESULTS:** Pathway-PRS for four mitochondrial pathways are significantly associated with increased AD risk: (i) response to oxidative stress (OR: 2.01 [95% Cl: 1.71, 2.37]); (ii) mitochondrial transport (OR: 1.81 [95% Cl: 1.55, 2.13]); (iii) hallmark oxidative phosphorylation (OR: 1.23 [95% Cl: 1.07, 1.41]); and (iv) mitochondrial membrane potential regulation (OR: 1.18 [95% Cl: 1.03, 1.36]).

**DISCUSSION:** Therapeutic approaches targeting these pathways may have potential for modifying AD pathogenesis. Further investigation is required to establish a causal role for these pathways in AD pathology.

## 1. Introduction

Alzheimer’s disease (AD) is a debilitating neurological condition characterized by memory deficits, cognitive and behavioural impairment [1] affecting more than 43.8 million people worldwide [2]. The classical neuropathological hallmarks of AD are the accumulation of amyloid-β peptides into extracellular neuritic plaques and hyperphosphorylated tau into intracellular neurofibrillary tangles in isocortical, subcortical and memory-associated regions of the brain [3]. The substantial attempts to develop drugs based on the role of amyloid-β and tau in AD pathogenesis have led to limited success in identifying disease modifying therapies [4]. This lack of success has led to the exploration of other potential causal mechanisms such as mitochondrial dysfunction.

Mitochondria are intracellular organelles involved in producing energy-carrying ATP molecules through oxidative phosphorylation (OXPHOS) and other cellular processes, including calcium homeostasis, response to oxidative stress (OXSTRESS) and apoptosis [5]. Each mitochondrion possesses its own ~16.5 kb circular genome (mtDNA) encoding 37 genes comprised of 2 ribosomal RNA genes, 22 tRNA genes, and 13 protein-coding genes. There are a further ~1,158 genes in the nuclear genome (nDNA) that also encode proteins involved in mitochondrial function, known as nuclear-encoded mitochondrial genes (nMT-genes) [6].

The mitochondrial cascade hypothesis of AD pathogenesis was first described in 2004 [7]. Briefly, baseline mitochondrial function is genetically determined and declines with age due to environmental and lifestyle factors [7]. This declining mitochondrial function is either the primary event initiating Aβ- or tau-induced toxicity (primary mitochondrial cascade) or a by-product of the amyloid cascade (secondary mitochondrial cascade) that results in AD pathology [8].

This hypothesis is supported by several lines of evidence. Early epidemiological studies reported a 3-9 fold higher AD risk associated with maternal AD history (possibly associated with maternally-inherited mtDNA) compared with paternal or no AD family history [9]. Altered mitochondrial structures and bioenergetics [10], reduced glucose utilization and functional deficits in several mitochondrial enzymes have been observed in AD brains ([11]). In transgenic *APP* mutant mice, upregulated compensatory mitochondrial mechanisms precede rather than follow amyloid-β plaque deposition and behavioural changes [12]. Cell culture studies demonstrate inhibition of mitochondrial COX enzyme activity [13], and increased mitochondrial-generated reactive oxygen species (ROS) [14] shift AβPP processing towards the amyloidogenic pathway.

Several mitochondrial pathways are dysregulated or dysfunctional in AD. ATP production is reduced due to OXPHOS dysfunction [15], mitochondrial transport is interrupted [16, 17], oxidative stress is increased [18], cellular apoptotic pathways are upregulated [19], intracellular neuronal calcium levels are increased and calcium buffering mechanisms are dysregulated [20], mitochondrial fission is increased and fusion decreased [21], mitophagy is defective [22], and mitochondrial membrane potential (mtΔΨ) is reduced [23]. However, the molecular mechanisms through which the mitochondria mediate, initiate or contribute to AD-related pathology remain unknown and highly debated.

Individually, most variants in the nuclear-encoded mitochondrial genome (nMT-DNA) have sub-threshold (p > 10^−8^) effects on AD risk [24] in genome-wide association studies (GWAS). Greater predictive power is obtained by investigating the combined effect of multiple SNPs as polygenic risk scores (PRS), which can be weighted by their GWAS effect sizes [25, 26]. PRSs can also be composed of genetic variants in multiple genes associated with the same biological pathway, forming a pathway-PRS. Taking this approach, we recently demonstrated that PRS composed of sub-threshold variants in nMT-genes is significantly associated with AD [27].

In this study, we use a pathway-based approach, constructing PRSs for sets of genes that encode components of mitochondrial pathways, to investigate their association with AD in a biologically informative way.

## 2. Methods

### 2.1 Alzheimer’s Disease Neuroimaging Initiative

This study used data from the Alzheimer’s Disease Neuroimaging Initiative (ADNI) [28], last accessed on 28 April 2019 (n = 2175). ADNI is a longitudinal study launched in 2004 with the objective of validating amyloid phenotyping, characterizing AD-associated biomarkers, and understanding the genetic underpinnings of AD to inform clinical trial design. ADNI’s AD diagnostic criteria are based on both clinical assessments and neurophysiological tests [28]. A case-control study design was employed based on diagnosis at last assessment for each participant.

Participants were excluded if they had any of the following: (i) missing diagnoses (n = 33), (ii) diagnosis of mild cognitive impairment but not AD (n = 558), (iii) missing *APOE* genotype (n = 44) or missing genome-wide sequencing data (n = 517). Only participants with self-reported ‘non-hispanic white’ ancestry were included in the study to avoid bias due to population stratification, resulting in exclusion of 169 additional samples. Final study sample included 854 participants.

### 2.2 Genotype data

Genotype data was obtained from the ADNI database (http://adni.loni.usc.edu). Details of the collection, curation, processing, and quality-control of ADNI data are described in detail elsewhere [28, 29]. Briefly, nDNA was extracted from whole blood and genotyped on Illumina GWAS arrays for ADNI1 (Illumina Human 610-Quad BeadChip), ADNI GO/2 participants (Illumina HumanOmniExpress BeadChip), and ADNI3 participants (Illumina Omni 2.5M) [28]. *APOE* genotyping of the two SNPs (rs429358, rs7412) that distinguish the ε2, ε3, and ε4 alleles was performed separately for all individuals, and quality controlled [29].

Standard GWAS quality control [30] performed on the ADNI genotype data included: removing variants with call rate <0.95, MAF <1%; deviation from Hardy-Weinberg equilibrium p<10-6 for CN and p<10-10 for AD cases; call rate <0.95; discordance between reported and genetically determined sex; cryptic sample relatedness (pi-hat threshold 0.2); and outlying heterozygosity. Genotype missingness < 5% was imputed using ADNI MAF alleles for PRS calculation. Principal component analysis (PCA) [31] was performed to correct for residual population stratification in the logistic regression model (see section 2.4).

### 2.3 Polygenic Risk Profiling

We constructed three whole-genome AD PRSs and 12 mitochondrial pathway-specific PRSs (Table 2) for each ADNI participant using the ‘standard weighted allele’ method implemented in PRSice2 and PRSet [32] for the whole genome and specific mitochondrial pathway gene sets, respectively.

SNPs were weighted by their GWAS effect sizes from the International Genomics of Alzheimer’s Project (IGAP) [24]. We retained GWAS SNPs with p-value threshold (P_T_) ≤ 0.5 for computing the PRS as this threshold provided the best model fit for our data (see Appendix C) and is supported by published evidence as the optimum threshold for estimating AD PRSs for common variants [26]. Linkage disequilibrium (LD) clumping was performed for the whole genome (250 kb window, r2 < 0.1) using PRSice-2 [32]. Set-based LD clumping was performed using PRSet [32] for pathway-specific polygenic risk scores to retain only data for SNPs in the gene-set regions (250 kb window, r2 < 0.1).

We omitted loci on sex chromosomes and in the major histocompatibility complex (MHC: 28.47 Mb–33.44 Mb, Chr6, GRCh37 [33]) because estimation of polygenic risk scores in these regions is difficult due to their genomic complexity, i.e. mismapping of reads due to high sequence homology between X and Y chromosomes, and high polymorphic diversity in the MHC region [25, 34]. As a result, 40 X-linked nMT-genes (Appendix A) and 5 nMT-genes in the MHC region (Appendix B) were excluded.

#### 2.3.1 Whole genome polygenic risk scores

Whole genome PRS was estimated in three ways: (i) Whole-genome, i.e., for all included SNPs; (ii) Excluding a 250 kb region of LD around the *APOE* gene (19:45409011 – 45412650 on GRCh37), to assess effects that are independent of the known AD-risk alleles of the *APOE* and *TOMM40* genes [35]; (iii) Excluding the *APOE* region and nMT-genes, to provide a baseline for estimating nMT-gene-specific effects.

#### 2.3.2 Mitochondrial pathway-specific polygenic risk scores

Mitochondrial pathway-specific PRSs were constructed for (i) 12 mitochondrial pathways represented by genesets obtained from the Molecular signatures database (MsigDB) [36] (Table-2) and (ii) the nMT-DNA geneset (comprising all nMT-genes) obtained from Mitocarta 2.0 [6]. Information about the curation and selection of these pathway genesets is detailed in Appendix D. These genesets include all nuclear genes that are known to influence or to be involved in mitochondrial function (Appendix E). Only the SNPs in introns and exons defined by GRCh37 gene boundaries [33] were included for PRS calculation.

Association of *TOMM40* with increased AD risk is confounded by its high LD with the *APOE* [37]. Therefore, for the nMT-DNA and mitochondrial transport genesets, two PRSs were calculated, one that included *TOMM40* and thus the confounding effect of *APOE*, and one that excluded *TOMM40* and thus excluding both the direct effect of *TOMM40* and the confounding effect of *APOE*. All polygenic risk scores were standardized to z-scores with respect to the sample mean.

### 2.4 Statistical Analysis

#### 2.4.1 Logistic Regression Modelling

Differences in the demographic characteristics of AD cases and CN controls were assessed via one-way ANOVA for continuous variables (age, years of education, Mini Mental State Examination (MMSE) score) and Fisher’s exact test for categorical variables (gender, *APOE* genotype). One-way ANOVA was also performed to assess if there were significant differences in the mean whole genome PRS and nMT-DNA PRS between the two diagnostic groups.

To evaluate the effect of the PRS on AD, a multivariable logistic regression model was run for AD cases vs CN controls. Age, sex, *APOE* ε4 allelic status (ε4 copy number = 0, 1, 2) and the first three principal components (representing population structure) were included as covariates.

All statistical analyses were performed in the R 3.4.4.

#### 2.4.2 Multiple testing burden correction

P-value significance was calculated after correcting for multiple testing burden using two methods (a) False Discovery rate (FDR < 0.05) using the Benjamini-Hochberg procedure [38] embedded within p.adjust() R function, and (b) competitive empirical p-value approach using PRSice and PRSet [32] to compare the outcome of adjusted p-value significance for each pathway using both methods.

A competitive p-value (*Competitive – P*) was obtained for each pathway-PRS as:

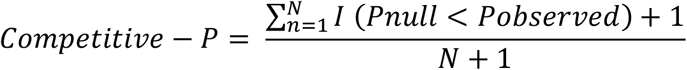

where *Pobserved* is the probability of the observed difference between cases and controls for each pathway-PRS, *Pnull* is obtained for SNPs randomly selected from the background exome in numbers (N) equivalent to those in the pathway-PRSs. Comparisons were made between pseudo-case and control groups to which individuals were randomly assigned. A *Pnull* distribution was obtained from 10,000 permutations.

## 3. Results

### 3.1 Research Cohort

Descriptive statistics for ADNI participants (n = 854; CN = 355, AD = 499) are presented in Table 1. There are significant differences between the CN and AD groups for years of education, *APOE* genotype, and MMSE score. Furthermore, the mean whole-genome PRS and complete nMT-DNA PRS are significantly different between the groups, with AD cases having a higher mean PRS compared to CN controls.

**Table 1.**
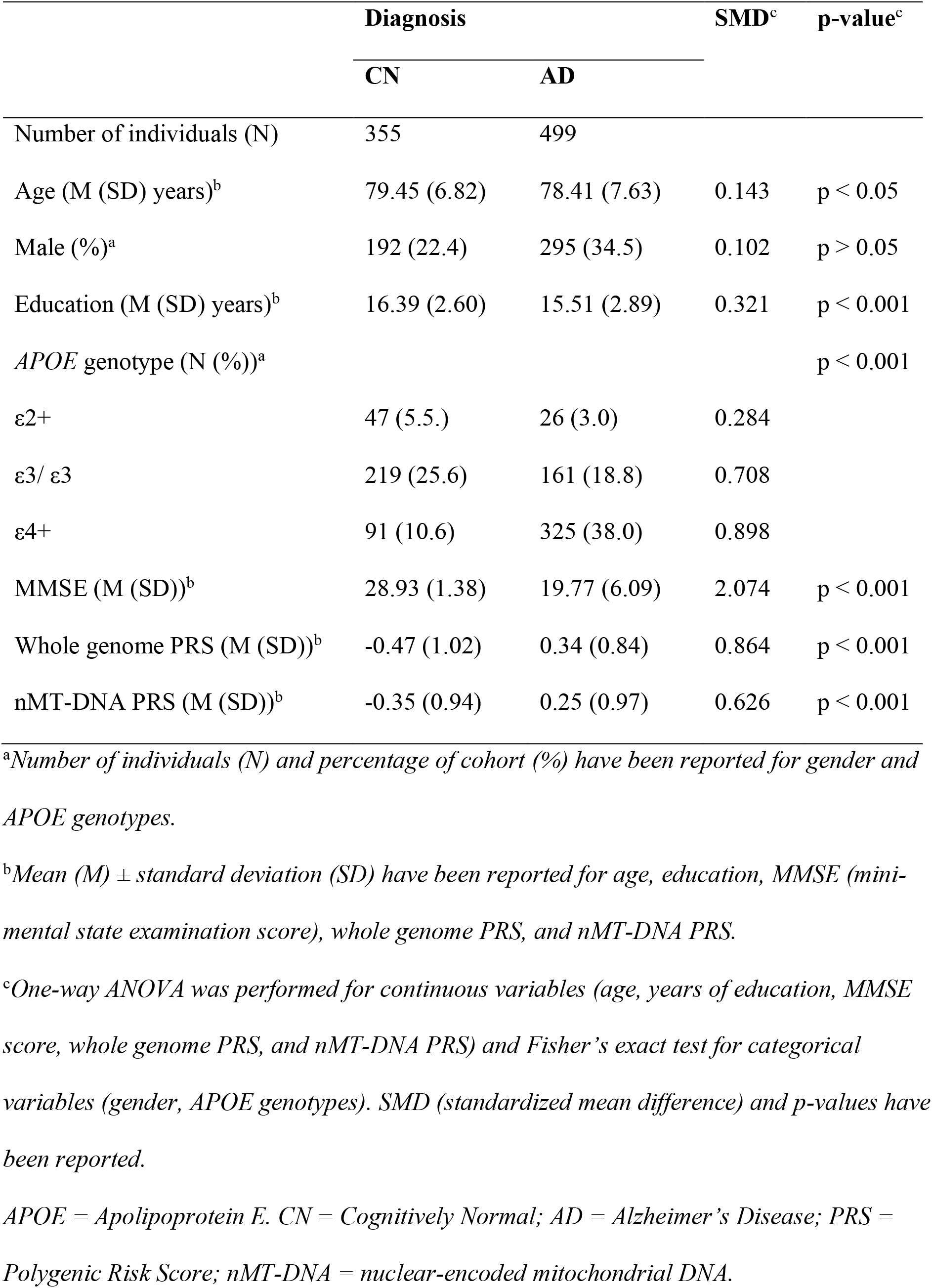
Demographics of ADNI participants (n = 854) included in this study.

|  | Diagnosis |  | SMD<sup>c</sup> | p-value<sup>c</sup> |
| --- | --- | --- | --- | --- |
|  | CN | AD |  |  |
| Number of individuals (N) | 355 | 499 |  |  |
| Age (M (SD) years)<sup>b</sup> | 79.45 (6.82) | 78.41 (7.63) | 0.143 | p < 0.05 |
| Male (%)<sup>a</sup> | 192 (22.4) | 295 (34.5) | 0.102 | p > 0.05 |
| Education (M (SD) years)<sup>b</sup> | 16.39 (2.60) | 15.51 (2.89) | 0.321 | p < 0.001 |
| *APOE* genotype (N (%))<sup>a</sup> |  |  |  | p < 0.001 |
| ε2+ | 47 (5.5.) | 26 (3.0) | 0.284 |  |
| ε3/ ε3 | 219 (25.6) | 161 (18.8) | 0.708 |  |
| ε4+ | 91 (10.6) | 325 (38.0) | 0.898 |  |
| MMSE (M (SD))<sup>b</sup> | 28.93 (1.38) | 19.77 (6.09) | 2.074 | p < 0.001 |
| Whole genome PRS (M (SD))<sup>b</sup> | -0.47 (1.02) | 0.34 (0.84) | 0.864 | p < 0.001 |
| nMT-DNA PRS (M (SD))<sup>b</sup> | -0.35 (0.94) | 0.25 (0.97) | 0.626 | p < 0.001 |

<sup>a</sup>Number of individuals (N) and percentage of cohort (%) have been reported for gender and *APOE* genotypes.
<sup>b</sup>Mean (M) ± standard deviation (SD) have been reported for age, education, MMSE (mini-mental state examination score), whole genome PRS, and nMT-DNA PRS.
<sup>c</sup>One-way ANOVA was performed for continuous variables (age, years of education, MMSE score, whole genome PRS, and nMT-DNA PRS) and Fisher's exact test for categorical variables (gender, *APOE* genotypes). SMD (standardized mean difference) and p-values have been reported.
*APOE* = Apolipoprotein E. CN = Cognitively Normal; AD = Alzheimer's Disease; PRS = Polygenic Risk Score; nMT-DNA = nuclear-encoded mitochondrial DNA.

### 3.2 Polygenic scores of the whole nuclear genome and AD risk

The whole-genome PRS is significantly associated with AD, with a 1 SD increase in the PRS associated with AD (OR: 7.42 [95% Cl: 5.62, 9.99]). These results remain highly significant even after (i) exclusion of *APOE* region ±250kb (OR: 6.37 [95% Cl: 4.81, 8.62]) and (ii) exclusion of both *APOE* region ±250kb and nMT-genes (OR: 6.40 [95% Cl: 4.83, 8.67]) (Fig 1). The OR decreases when these gene regions are excluded, but the confidence intervals largely overlap (Fig 1). These ORs are substantially greater than the OR for *APOE* ±250kb region alone (OR: 1.98 [95% Cl: 1.68, 2.35).

**Fig 1.**
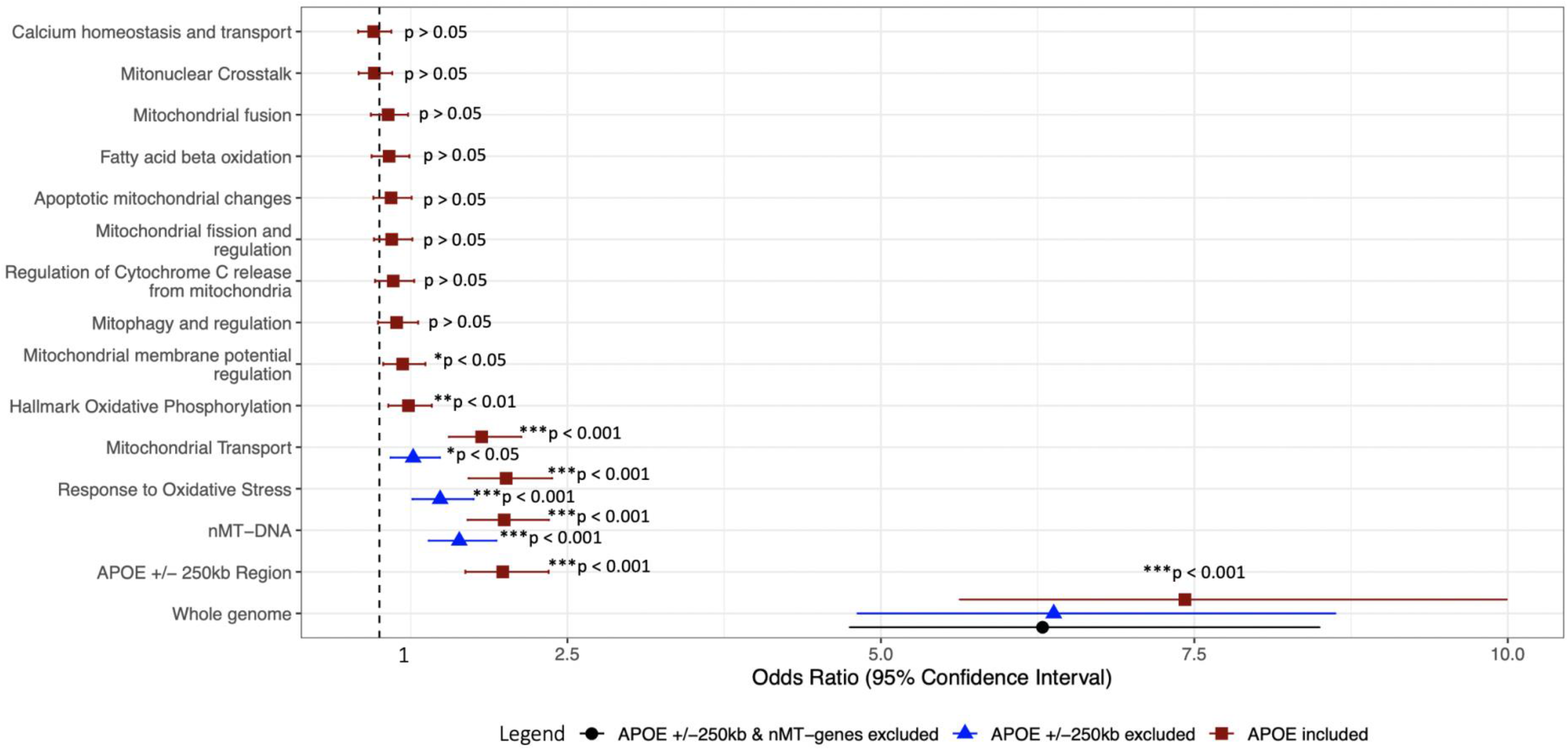
Odds Ratio estimates (95% Confidence intervals) for polygenic risk scores regressed with AD diagnosis. Significance of FDR-adjusted p-values reported here, are interpreted as follows: * p < 0.05 (significant); ** p < 0.01 (very significant); *** p < 0.001 (highly significant). Red squares denote inclusion of variants in the APOE region, blue triangles denote exclusion of the APOE ± 250 kb region, and the black circle denotes exclusion of the APOE ± 250 kb region and the complete nuclear-encoded mitochondrial genome (nMT-DNA).

### 3.3 Polygenic scores of nMT-DNA and mitochondrial pathways and AD risk

The nMT-DNA PRS is significantly associated with AD (OR: 1.99 [95% Cl: 1.70, 2.35]). In the pathway analyses, four mitochondrial pathways are significantly associated with AD: (i) OXSTRESS (OR: 2.01 [95% Cl: 1.71, 2.37]); (ii) mitochondrial transport (OR: 1.81 [95% Cl: 1.55, 2.13]); (iii) oxidative phosphorylation (OR: 1.23 [95% Cl: 1.07, 1.41]); and (iv) mtΔΨ regulation (OR: 1.18 [95% Cl: 1.03, 1.36]; Fig-1). For the mitophagy and regulation pathway-PRS (OR: 1.13 [95% Cl: 0.98, 1.31]), the FDR-adjusted p-value is non-significant (P_FDR_ > 0.05), however the competitive p-value is significant (P_competitive_ < 0.05).

The associations of nMT-DNA PRS (OR: 1.63 [95% Cl: 1.37, 1.94]) and OXSTRESS (OR: 1.57 [95% Cl: 1.32, 1.88]) pathways remain significant after exclusion of *APOE* ±250kb region (Table 3). For the Mitochondrial transport pathway-PRS (OR: 1.23 [95% Cl: 1.03, 1.48]), the FDR-adjusted p-value remains significant (P_FDR_ < 0.05), but the competitive p-value becomes non-significant (P_competitive_ > 0.05) when the *APOE* ±250kb region is excluded. Omission of the *APOE* ±250kb region was not necessary for the other pathways because they do not contain genes within this region.

**Table 2.** Genomic regions and pathway gensets for which polygenic risk scores were calculated.

| **Polygenic Risk Score** | **Total genes<sup>a</sup>** | **nMT-genes<sup>a</sup>** | **SNPs<sup>b</sup>** |
| --- | --- | --- | --- |
| Whole nuclear genome | N/A | N/A | 515164 |
| Whole nuclear genome (excluding *APOE* ± 250 kb) | N/A | N/A | 515058 |
| Whole nuclear genome (excluding *APOE* ±250kb and nMT-DNA<sup>b</sup>) | N/A | N/A | 83066 |
| nMT-DNA<sup>c</sup> | 1158 | 1158 | 13992 |
| Response to Oxidative Stress | 352 | 60 | 5803 |
| Hallmark Oxidative Phosphorylation | 245 | 211 | 2399 |
| Mitochondrial Transport | 179 | 107 | 2394 |
| Apoptotic mitochondrial changes | 58 | 25 | 922 |
| Mitochondrial membrane potential regulation | 54 | 21 | 1124 |
| Mitonuclear crosstalk | 38 | 0 | 1061 |
| Mitochondrial fission and regulation | 24 | 13 | 652 |
| Fatty acid beta-oxidation | 21 | 14 | 211 |
| Calcium homeostasis and transport | 19 | 7 | 402 |
| Mitochondrial fusion | 19 | 11 | 262 |
| Mitophagy and regulation | 171 | 24 | 3401 |

<sup>a</sup>Total number of nuclear genes and nuclear-encoded mitochondrial genes (nMT-genes) in each geneset have been reported.
<sup>b</sup>Total number of SNPs included for polygenic risk score calculation for each pathway are reported.
<sup>c</sup>nMT-DNA comprises of the complete geneset of nuclear-encoded mitochondrial genome.
*APOE* = Apolipoprotein E.

**Table 3.** Significant associations of polygenic risk scores for different genomic regions or pathways with Alzheimer’s Disease.

| Genomic region/ pathway | Including *APOE* region | | | | Excluding *APOE* ($\pm 250$ kb window) | | | |
| --- | --- | --- | --- | --- | --- | --- | --- | --- |
|  | Beta<sup>a</sup> | SE<sup>b</sup> | FDR-adjusted | Competitive | Beta<sup>a</sup> | SE<sup>b</sup> | FDR-adjusted | Competitive |
|  |  |  | p-value<sup>b</sup> | p-value |  |  | p-value<sup>b</sup> | p-value |
| Whole genome | 13.70 | 0.14 | $1.36 \times 10^{-41}$ | $1.07 \times 10^{-65}$ | 12.46 | 0.14 | $9.78 \times 10^{-35}$ | $1.34 \times 10^{-52}$ |
| *APOE* +/- 250 kb region | 8.08 | 0.08 | $2.33 \times 10^{-15}$ | $9.98 \times 10^{-19}$ | 8.08 | 0.08 | N/A | N/A |
| nMT-DNA | 8.38 | 0.08 | $3.68 \times 10^{-16}$ | $1.23 \times 10^{-15}$ | 5.63 | 0.08 | $1.65 \times 10^{-8}$ | $5.09 \times 10^{-7}$ |
| Response to oxidative stress | 8.32 | 0.08 | $4.37 \times 10^{-16}$ | $2.68 \times 10^{-17}$ | 5.01 | 0.09 | $7.51 \times 10^{-6}$ | $9.02 \times 10^{-6}$ |
| Mitochondrial transport | 7.40 | 0.08 | $3.97 \times 10^{-13}$ | $4.75 \times 10^{-12}$ | 2.27 | 0.09 | $7.12 \times 10^{-3}$ | $6.50 \times 10^{-2}$ |
| Hallmark oxidative phosphorylation | 2.90 | 0.07 | $9.07 \times 10^{-3}$ | $2.41 \times 10^{-3}$ | 2.39 | 0.07 | N/A | N/A |
| Mitochondrial membrane potential regulation | 2.37 | 0.07 | $3.78 \times 10^{-2}$ | $1.85 \times 10^{-2}$ | 2.83 | 0.07 | N/A | N/A |
| Mitophagy and regulation | 1.80 | 0.07 | $1.34 \times 10^{-1}$ | $1.72 \times 10^{-2}$ | 1.87 | 0.07 | N/A | N/A |
| Regulation of cytochrome C release from mitochondria | 1.46 | 0.07 | $2.38 \times 10^{-1}$ | $5.49 \times 10^{-2}$ | 2.19 | 0.07 | N/A | N/A |
| Mitochondrial fission and regulation | 1.32 | 0.07 | $2.77 \times 10^{-1}$ | $9.13 \times 10^{-2}$ | 1.33 | 0.07 | N/A | N/A |
| Apoptotic mitochondrial changes | 1.26 | 0.07 | $2.82 \times 10^{-1}$ | $3.50 \times 10^{-1}$ | 2.10 | 0.07 | N/A | N/A |
| Mitochondrial fusion | 0.96 | 0.07 | $3.88 \times 10^{-1}$ | $6.41 \times 10^{-1}$ | 1.16 | 0.07 | N/A | N/A |
| Fatty acid beta oxidation | 1.05 | 0.07 | $3.66 \times 10^{-1}$ | $4.25 \times 10^{-1}$ | 1.73 | 0.07 | N/A | N/A |
| Calcium homeostasis and transport | -0.66 | 0.07 | $5.44 \times 10^{-1}$ | $9.28 \times 10^{-1}$ | -0.73 | 0.07 | N/A | N/A |
| Mitonuclear crosstalk | -0.58 | 0.07 | $5.60 \times 10^{-1}$ | $4.78 \times 10^{-1}$ | -0.28 | 0.07 | N/A | N/A |

<sup>a</sup>Beta denotes beta estimates from multivariate regression between pathway-PRS and diagnosis.
<sup>b</sup>Standard errors (SE) and competitive empirical p-value of association are reported.
APOE = Apolipoprotein E. nMT-DNA represents the complete nuclear-encoded mitochondrial geneset.

The phenotypic variance explained by the genetic contribution of each mitochondrial pathway is presented in Appendix F.

## 4. Discussion

In this study, we investigated the association of AD polygenic risk scores composed of genetic variants located within genes associated with known mitochondrial pathways with AD risk. We found that pathway-PRS composed of the complete nuclear-encoded mitochondrial genome and genes involved in (i) response to oxidative stress, (ii) mitochondrial transport, (iii) hallmark oxidative phosphorylation, and (iv) mitochondrial membrane potential regulation were associated with increased AD risk. The results obtained by the pathway-based approach used here suggest that SNPs in nMT-genes and other nuclear genes involved in mitochondrial pathways significantly contribute to AD risk, suggesting therapeutic potential in targeting them.

Previous multi-omics studies support the role of mitochondrial pathways in AD. For instance, Mostafavi et al. (2018) built molecular networks using modules of co-expressed genes associated with AD and its endophenotypes. They found three modules enriched for gene ontology categories related to mitochondria showing a positive correlation with histopathological β-amyloid burden, cognitive decline, and clinical diagnosis of AD [39]. Similarly, Johnson et al. (2020) conducted co-expression network analysis of AD brains and found that protein co-expression families involved in mitochondrial metabolism strongly correlated with AD, and showed the strongest differences by case status [40]. Muraoka et. Al (2020) performed proteomic profiling of AD brain tissues and found that 148 proteins unique to AD group belonged to gene ontology categories associated with mitochondrial metabolism [41].

Our results add to accumulating evidence from genetic, clinical, model cell and animal studies supporting the involvement of these mitochondrial pathways and nMT-genes in AD. We found that a 1 SD increase in the OXPHOS pathway-PRS is associated with 1.23 times increased likelihood of developing AD. The importance of OXPHOS pathway in generating ATP-energy to support the high cellular energy demand in the brain and the body is well-established [5]. Reduced ATP production due to mitochondrial OXPHOS dysfunction activates a cascade of events leading to neural cell death observed in AD-associated neurodegeneration [15]. Importantly, dysregulation of gene-networks and gene-regulation can also facilitate OXPHOS dysfunction, as demonstrated for *PTCD1* using knock-out and cell-culture models [42, 43].

We found that a 1 SD increase in the mitochondrial transport pathway-PRS is associated with 1.81 times increased likelihood of developing AD. Disruption of mitochondrial transport can lead to defective communication with the nucleus and other cytosolic components. Current evidence indicates that defective mitochondrial transport in AD is primarily due to Aβ- and tau-interactions blocking mitochondrial channels such as TOMM40 [16], TIM23 [16], TOMM22 [44], and VDAC1 [17]. Hence, it is possible that mitochondrial transport dysfunction is a secondary effect of Aβ and tau toxicity. The contribution of *TOMM40* to PRS for the mitochondrial transport geneset is difficult to delineate due to confounding from its high LD with *APOE* [37].

We found that a 1SD increase in the OXTRESS pathway-PRS is associated with 2.01 times increased likelihood of developing AD. The OXTRESS pathway, which, when upregulated, activates the eIF2α/ATF4 axis increasing expression of stress response genes [45], is strongly associated with AD pathology and causes calcium dyshomeostasis, loss of mtΔΨ, high mutation rates, interrupted gene transcription and regulation due to high ROS-mediated cellular damage [18]. The OXSTRESS geneset contains the *APOE* gene. Increasing evidence suggests that *APOE* genotype influences mitochondrial stress-related processes in an APOE isoform-specific manner [46]. Therefore, it is likely that SNPs from the *APOE* region contribute to the higher pathway-PRS and AD association.

Finally, we found that a 1 SD increase in the mtΔΨ regulation pathway-PRS is associated with 1.18 times increased likelihood of developing AD. This pathway contains genes encoding proton pumps that regulate the electric charge potential across the mitochondrial membrane to maintain mitochondrial homeostasis and drive processes like the respiratory chain [47]. There is evidence for the formation of a complex through interaction of Aβ with cyclophilin D (CypD) that provokes mitochondrial and neuronal perturbation, as observed in transgenic *APP* mutant mice [48]. This leads to the Ca^2+^-dependent formation and opening of the mitochondrial membrane permeability transition pore (mPTP), which results in decreased mitochondrial membrane potential, and is accompanied by increased oxidative stress, compromised OXPHOS, release of cytochrome, and impaired axonal mitochondrial transport [49]. Thus, mPTP and CypD have been proposed as potential drug targets. Recent evidence showed that pharmacological blockage of mPTP with Cyclosporine A (CsA) significantly improved mitochondrial and cytosolic calcium dysregulation in AD fibroblasts [23]. Future research is required to investigate their therapeutic potential in AD brain models.

Moderate to high heritability of late-onset AD (LOAD) is indicated by SNP (~ 53%) [50] and twin studies (~ 60-80%) [51]. *APOE* is the strongest known genetic predictor of LOAD risk. It explains up to 13% of the phenotypic variance [50], which implies that at least 37% of the phenotypic variance is explained by genetic variation outside this region [50]. This could explain our result that the OR estimates for AD risk remain significant regardless of the inclusion and exclusion of *APOE* region (±250 kb window) from the whole genome PRS (Fig-1).

The nMT-DNA, mitochondrial transport, and OXSTRESS genesets contain genes located in the *APOE* ±250 kb region [52]. Interestingly, the OR estimates for nMT-DNA and OXSTRESS pathways are similar to the estimate for the APOE ±250kb region. Yet, the association of nMT-DNA and OXSTRESS pathway-PRS with AD remains significant even after the exclusion of the *APOE* ±250 kb region. Thus, reinforcing the contribution of non-*APOE* genes in conferring substantial polygenic risk for these pathways. However, the competitive p-value for mitochondrial transport is non-significant on the exclusion of *APOE* ±250 kb region, which includes the *TOMM40* gene. This result may reflect the known high risk of *APOE/TOMM40* loci [37]. Because of high LD, the contributions of these two loci to PRS cannot be separated. Exclusion of the region may, therefore, result from either removal of a real effect of *TOMM40* variation or removal of a confounding effect of *APOE* variation.

These results should be interpreted in conjunction with some study limitations. First, the ADNI cohort is relatively small and our results require replication. Second, only participants of European ancestry were included, therefore these results may not be generalizable to other ancestrally diverse populations. Finally, our pathway-PRS may underestimate of true genetic risk conferred by mitochondrial pathways. We could not include mtDNA SNPs previously implicated with AD risk [53] since large-scale AD GWAS for mtDNA variants are currently unavailable.

The primary strength of this study is the pathway-driven, biologically informed approach to polygenic risk scoring. The pathway-PRS OR estimates presented in this paper are comparable to those previously reported for high risk single GWS variants [52]. This is because pathway-PRS effectively capture the cumulative small-effects of sub-threshold variants within genes involved in mitochondrial pathways. This results in a larger combined effect size and hence higher statistical power to detect association with AD than association testing of GWS variants individually.

In conclusion, this study demonstrated that the genetic variation within the OXPHOS, mitochondrial transport, OXSTRESS, and mtΔΨ regulation pathways captured by pathway-PRS significantly influences AD risk. These findings contribute to the growing evidence of a mitochondrial role in AD and suggest these pathways as potential targets in ameliorating AD pathogenesis. However, it remains to be determined if mitochondrial dysfunction is the primary cause of AD pathogenesis. Further investigations are required to validate and establish the causal role of nMT-genes and pathways in AD pathology.

## Abbreviations

nMT-DNA: nuclear-encoded mitochondrial genome
nMT-genes: nuclear-encoded mitochondrial genes
OXPHOS: Oxidative phosphorylation
OXSTRESS: Response to oxidative stress
PRS: Polygenic risk scores
Pathway-PRS: Pathway polygenic risk scores

## Acknowledgements

Data for this project was made available via the Alzheimer’s Disease Neuroimaging Initiative (ADNI). ADNI is funded by the United States (National Institutes of Health, United States Grant U01 AG024904) and DOD ADNI (Department of Defense, United States award number W81XWH-12-2-0012). ADNI is funded by the National Institute on Aging, United States, the National Institute of Biomedical Imaging and Bioengineering, and through generous contributions from the following: AbbVie; Alzheimer’s Association; Alzheimer’s Drug Discovery Foundation; Araclon Biotech; BioClinica, Inc; Biogen; Bristol-Myers Squibb Company; CereSpir, Inc; Cogstate; Eisai Inc; Elan Pharmaceuticals, Inc; Eli Lilly and Company; EuroImmun; F. Hoffmann-La Roche Ltd and its affiliated company Genentech, Inc; Fujirebio; GE Healthcare; IXICO Ltd; Janssen Alzheimer Immunotherapy Research & Development, LLC; Johnson & Johnson Pharmaceutical Research & Development LLC; Lumosity; Lundbeck; Merck & Co, Inc; Meso Scale Diagnostics, LLC; NeuroRx Research; Neurotrack Technologies; Novartis Pharmaceuticals Corporation; Pfizer Inc; Piramal Imaging; Servier; Takeda Pharmaceutical Company; and Transition Therapeutics. The Canadian Institutes of Health Research is providing funds to support ADNI clinical sites in Canada. Private sector contributions are facilitated by the Foundation for the National Institutes of Health (www.fnih.org). The grantee organization is the Northern California Institute for Research and Education, and the study is coordinated by the Alzheimer’s Therapeutic Research Institute at the University of Southern California. ADNI data are disseminated by the Laboratory for Neuro Imaging at the University of Southern California.

## Appendix

Appendices A-F provided.

