## Supplemental file for "Mitochondrial pathway polygenic risk scores are associated with Alzheimer’s Disease"

### Appendices

### A. Nuclear-encoded mitochondrial genes on Chromosome X

**Table A.1 Nuclear encoded mitochondrial genes on Chromosome X excluded from the ADNI genotyping data for polygenic score calculation.**

| **Ensembl ID** | **Symbol** | **Description** | **Chr Start** | **Chr End** |
|---|---|---|---|---|
| ENSG00000131828 | *TST* | thiosulfate sulfurtransferase (rhodanese) | 19362010 | 19379825 |
| ENSG00000156709 | *TSTD1* | thiosulfate sulfurtransferase (rhodanese)-like domain containing 1 | 129263337 | 129299861 |
| ENSG00000155008 | *TSTD3* | thiosulfate sulfurtransferase (rhodanese)-like domain containing 3 | 84258897 | 84348323 |
| ENSG00000067829 | *TTC19* | tetratricopeptide repeat domain 19 | 153051220 | 153059978 |
| ENSG00000004961 | *TUBB3* | tubulin, beta 3 class III | 11129405 | 11141204 |
| ENSG00000131269 | *TUFM* | Tu translation elongation factor, mitochondrial | 74273006 | 74376175 |
| ENSG00000005022 | *TXN2* | thioredoxin 2 | 118602362 | 118605359 |
| ENSG00000067992 | *TXNDC12* | thioredoxin domain containing 12 (endoplasmic reticulum) | 24483343 | 24568583 |
| ENSG00000147123 | *TXNRD1* | thioredoxin reductase 1 | 47001614 | 47004609 |
| thioredoxin reductase 2 | *TXNRD2* | 40482817 40483391 | | |
| ENSG00000126768 | *TYSND1* | trypsin domain containing 1 | 48750729 | 48755426 |
| ENSG00000198814 | *UCP1* | uncoupling protein 1 (mitochondrial, proton carrier) | 30671475 | 30749577 |
| ENSG00000072506 | *UCP2* | uncoupling protein 2 (mitochondrial, proton carrier) | 53458205 | 53461323 |
| ENSG00000126953 | *UCP3* | uncoupling protein 3 (mitochondrial, proton carrier) | 100600643 | 100603957 |
| ENSG00000131174 | *UNG* | uracil-DNA glycosylase | 77154960 | 77160881 |
| ENSG00000188917 | *UQCC1* | ubiquinol-cytochrome c reductase complex assembly factor 1 | 100264333 | 100307105 |
| ENSG00000169239 | *UQCC2* | ubiquinol-cytochrome c reductase complex assembly factor 2 | 15756411 | 15805748 |
| ENSG00000158578 | *UQCR10* | ubiquinol-cytochrome c reductase, complex III subunit X | 55035487 | 55057497 |
| ENSG00000165775 | *UQCR11* | ubiquinol-cytochrome c reductase, complex III subunit XI | 154255063 | 154285191 |
| ENSG00000184831 | *UQCRB* | ubiquinol-cytochrome c reductase binding protein | 23851464 | 23926057 |
| ENSG00000102078 | *UQCRC1* | ubiquinol-cytochrome c reductase core protein I | 129473861 | 129507335 |
| ENSG00000125356 | *UQCRC2* | ubiquinol-cytochrome c reductase core protein II | 119005733 | 119010629 |
| ENSG00000123130 | *UQCRFS1* | ubiquinol-cytochrome c reductase, Rieske iron-sulfur polypeptide 1 | 23721776 | 23761407 |
| ENSG00000169084 | *UQCRH* | ubiquinol-cytochrome c reductase hinge protein | 2137554 | 2419015 |

| | | | | |
|---|---|---|---|---|
| ENSG00000176274 | *UQCRQ* | ubiquinol-cytochrome c reductase, complex III subunit VII, 9.5kDa | 103343897 | 103401708 |
| ENSG00000069509 | *USMG5* | up-regulated during skeletal muscle growth 5 homolog (mouse) | 44382884 | 44402221 |
| ENSG00000036473 | *VARS2* | valyl-tRNA synthetase 2, mitochondrial | 38211735 | 38280703 |
| ENSG00000077713 | *VDAC1* | voltage-dependent anion channel 1 | 118533257 | 118588437 |
| ENSG00000101986 | *VDAC2* | voltage-dependent anion channel 2 | 152990322 | 153010216 |
| ENSG00000182712 | *VDAC3* | voltage-dependent anion channel 3 | 154289899 | 154299547 |
| ENSG00000069535 | *VWA8* | von Willebrand factor A domain containing 8 | 43625856 | 43741721 |
| ENSG00000178605 | *WARS2* | tryptophanyl tRNA synthetase 2, mitochondrial | 221425 | 230887 |
| ENSG00000169100 | *WBSCR16* | Williams-Beuren syndrome chromosome region 16 | 1505044 | 1511039 |
| ENSG00000189221 | *WDR81* | WD repeat domain 81 | 43514154 | 43606071 |
| ENSG00000185973 | *XPNPEP3* | X-prolyl aminopeptidase (aminopeptidase P) 3, putative | 154718672 | 154842622 |
| ENSG00000123131 | *XRCC6BP1* | XRCC6 binding protein 1 | 23685644 | 23704514 |
| ENSG00000169188 | *YARS2* | tyrosyl-tRNA synthetase 2, mitochondrial | 55026755 | 55034306 |
| ENSG00000068366 | *YBEY* | ybeY metallopeptidase (putative) | 108884563 | 108976621 |
| ENSG00000174740 | *YME1L1* | YME1-like 1 ATPase | 90689596 | 90693583 |
| ENSG00000214827 | *ZADH2* | zinc binding alcohol dehydrogenase domain containing 2 | 154292308 | 154299547 |

### B. Nuclear-encoded mitochondrial genes on Major Histocompatibility Complex

**Table B.1 Nuclear-encoded mitochondrial genes on Major histocompatibility complex (28.47 Mb – 33.44 Mb on chromosome 6, GRCh37 coordinates) excluded from the ADNI genotyping data for polygenic score calculation.**

| Ensembl ID | Symbol | Description | Chr Start | Chr End |
|---|---|---|---|---|
| ENSG00000172171 | *HSD17B8* | Hydroxysteroid (17-beta) dehydrogenase 8 | 33172413 | 33174608 |
| ENSG00000158042 | *C6orf136* | chromosome 6 open reading frame 136 | 30614815 | 30620987 |
| ENSG00000036473 | *VARS2* | valyl-tRNA synthetase 2, mitochondrial | 30881981 | 30894235 |
| ENSG00000115317 | *MRPS18B* | mitochondrial ribosomal protein S18B | 30585485 | 30594174 |
| ENSG00000198721 | *RPS18* | ribosomal protein S18 | 33239851 | 33244281 |

### C. P-value thresholding

P-value thresholding was performed to retain more informative SNPs and reduce background noise. The best p-value threshold was determined by calculating PRS across a range of thresholds and comparing their model fit for (i) the complete nuclear genome and (ii) nuclear-encoded mitochondrial genome (nMT-genes with +/-10kb window). This model fit was calculated by subtracting the Nagelkerke's $r^2$ variance of the full regression model (consisting of both polygenic score and covariates) from the $r^2$ variance null model (consisting of covariates only). Here, covariates include the number of *APOE ε4* alleles, age, sex, and the first three principal components.

Eq. (C.1) Calculation of model-fit for a p-value threshold

$$\begin{aligned} Model\ fit\ R^2 = {} & R^2\ of\ full\ model\ (Phenotype \sim PRS + covariates) \\ & - R^2\ of\ Null\ model\ (Phenotype \sim covariates) \end{aligned}$$

The GWAS association p-value threshold of 0.5 provides a high model fit for our data. Thus, SNPs with GWAS p-value of association ≤ 0.5 were used for PRS calculations for all genomic regions and pathways to maintain consistency.

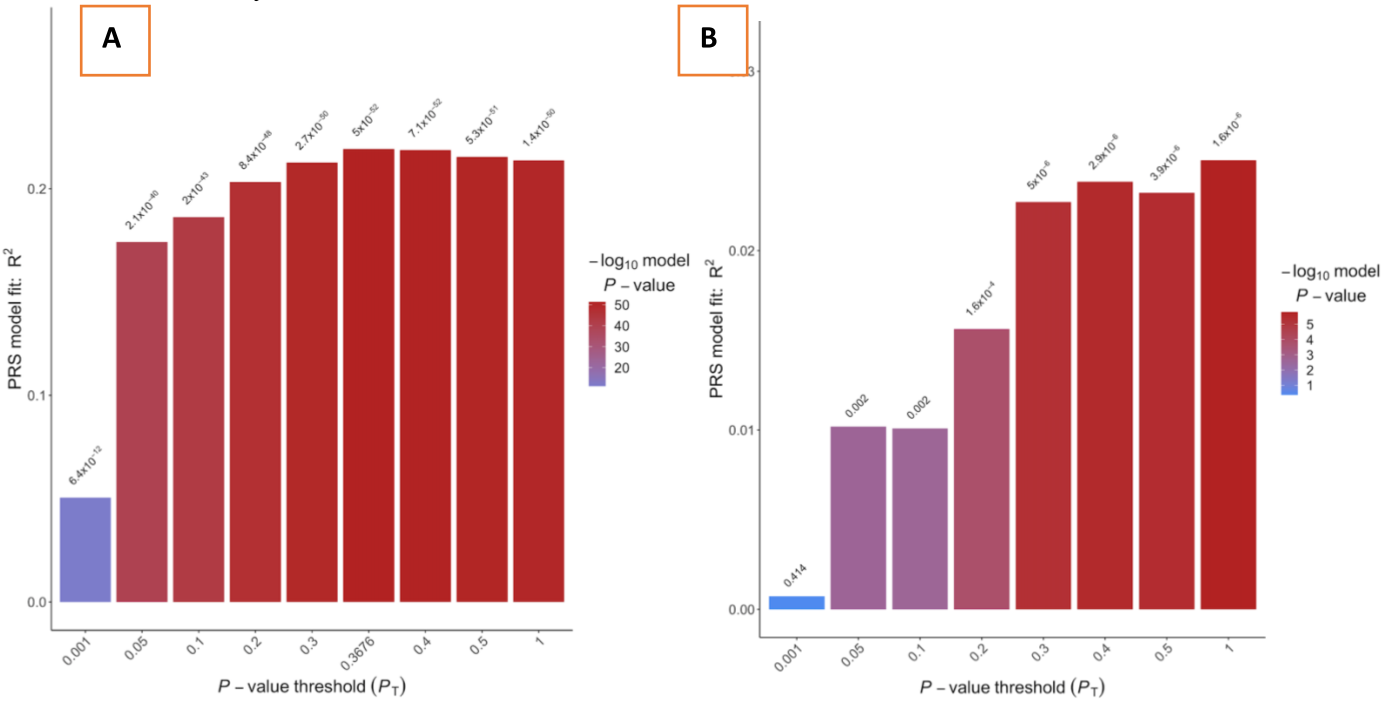

***Fig. C.1 Model fit for polygenic score calculations across a range of thresholds*** *for the (A) whole nuclear genome and (B) complete nuclear-encoded mitochondrial genome.*

### D. Curation of Mitochondrial Pathway Genesets

Downloaded and combined the following mitochondrial pathway genesets from MsigDB database (http://software.broadinstitute.org/gsea/msigdb/index.jsp) on 8 April 2019.

**Table D.1 List of MsigDB gene sets combined to form a pathway geneset.**

| Final Pathway Geneset | MsigDB genesets combined | No. of Genes |
|---|---|---|
| nMT-DNA[a] | • All mitochondrial genes list from Human MitoCarta2.0[a] | 1158 |
| Hallmark oxidative phosphorylation | • Hallmark Oxidative Phosphorylation<br>• Mitochondrial Respiratory Chain<br>• GO Mitochondrial Respiratory Chain Complex Assembly | 245 |
| Mitochondrial Transport | • Mitochondrial Transport<br>• GO Mitochondrial Transport | 179 |
| Response to Oxidative Stress | • GO Response to Oxidative stress | 352 |
| Apoptotic Mitochondrial Changes | • GO Apoptotic Mitochondrial Changes<br>• Apoptotic Mitochondrial Changes | 58 |
| Calcium Homeostasis and Transport | • GO Mitochondrial Calcium Homeostasis<br>• GO Mitochondrial Calcium Transport | 19 |
| Mitochondrial Fission | • GO Mitochondrial Fission<br>• GO Positive regulation of mitochondrial fission | 24 |
| Mitochondrial Fusion | • GO Mitochondrial Fusion | 19 |
| Mitophagy and regulation | • GO Macromitophagy<br>• GO regulation of mitophagy | 171 |
| Mitonuclear Crosstalk[b] | • Anterogade Signalling[b]<br>• Reterogade signalling[b] | 38 |
| Fatty Acid Beta oxidation | • Reactome Mitochondrial Fatty Acid Beta Oxidation | 21 |
| Mitochondrial Membrane Potential regulation | • GO Regulation of Mitochondrial membrane potential | 54 |

[a]nMT-DNA comprised of all nuclear-encoded mitochondrial genes, geneset obtained from [1].
[b]Mitonuclear Crosstalk geneset was obtained from [2].

### E. Genes included in mitochondrial pathway genesets

**Table E.1 List of all genes included in each mitochondrial pathway geneset.**

| **Hallmark oxidative phosphorylation & Respiratory Chain** |
|---|
| *ABCB7 ACAA1 ACAA2 ACADM ACADSB ACADVL ACAT1 ACO2 AFG3L2 AIFM1 ALAS1 ALDH6A1 ATP1B1 ATP5A1 ATP5B ATP5C1 ATP5D ATP5E ATP5F1 ATP5G1 ATP5G2 ATP5G3 ATP5H ATP5I ATP5J ATP5J2 ATP5L ATP5O ATP6AP1 ATP6V0B ATP6V0C ATP6V0E1 ATP6V1C1 ATP6V1D ATP6V1E1 ATP6V1F ATP6V1G1 ATP6V1H BAX BCKDHA BDH2 BRP44L CASP7 COX10 COX11 COX15 COX17 COX4I1 COX5A COX5B COX6A1 COX6B1 COX6C COX7A2 COX7A2L COX7B COX7C COX8A CPT1A CS CYB5A CYB5R3 CYC1 CYCS DECR1 DLAT DLD DLST ECH1 ECHS1 ECI1 ETFA ETFB ETFDH FDX1 FH FXN GLUD1 GOT2 GPI GPX4 GRPEL1 HADHA HADHB HCCS HSD17B10 HSPA9 HTRA2 IDH1 IDH2 IDH3A IDH3B IDH3G IMMT ISCA1 ISCU LDHA LDHB LRPPRC MAOB MDH1 MDH2 MFN2 MGST3 MRPL11 MRPL15 MRPL34 MRPL35 MRPS11 MRPS12 MRPS15 MRPS22 MRPS30 MTRF1 MTRR MTX2 NDUFA1 NDUFA2 NDUFA3 NDUFA4 NDUFA5 NDUFA6 NDUFA7 NDUFA8 NDUFA9 NDUFAB1 NDUFB1 NDUFB2 NDUFB3 NDUFB4 NDUFB5 NDUFB6 NDUFB7 NDUFB8 NDUFC1 NDUFC2 NDUFS1 NDUFS2 NDUFS3 NDUFS4 NDUFS6 NDUFS7 NDUFS8 NDUFV1 NDUFV2 NNT NQO2 OAT OGDH OPA1 OXA1L PDHA1 PDHB PDHX PDK4 PDP1 PHB2 PHYH PMPCA POLR2F POR PRDX3 RETSAT RHOT1 RHOT2 SDHA SDHB SDHC SDHD SLC25A11 SLC25A12 SLC25A20 SLC25A3 SLC25A4 SLC25A5 SLC25A6 SUCLA2 SUCLG1 SUPV3L1 SURF1 TCIRG1 TIMM10 TIMM13 TIMM17A TIMM50 TIMM8B TIMM9 TOMM22 TOMM70A UQCR10 UQCR11 UQCRB UQCRC1 UQCRC2 UQCRFS1 UQCRH UQCRQ VDAC1 VDAC2 VDAC3 BCS1L ND1 NDUFA13 AARS2 ACAD9 C11orf83 C12orf62 C18orf55 C19orf79 C20orf7 C2orf56 C6orf125 C7orf44 C8orf38 CCDC56 CHCHD5 CHCHD8 COX19 ECSIT FAM36A FOXRED1 IMMP1L IMMP2L NDUFA10 NDUFA11 NDUFA12 NDUFAF1 NDUFAF2 NDUFAF3 NDUFAF4 NDUFB10 NDUFB11 NDUFB9 NDUFS5 NDUFV3 NUBPL SAMM50 SDHAF1 SLC25A33 TAZ TFAM TIMMDC1 TMEM126B TTC19 UQCC* |
| **Mitochondrial Transport** |
| *ACAA2 ACACA ACACB AFG3L2 AIP ALKBH7 ATF2 ATP5A1 ATP5B ATP5C1 ATP5D ATP5E ATP5F1 ATP5G1 ATP5G2 ATP5G3 ATP5H ATP5I ATP5J ATP5J2 ATP5L ATP5O ATP5S ATPIF1 BAD BAK1 BAX BBC3 BCL2 BCL2L1 BCL2L11 BHLHA15 BID BLOC1S2 BMF BNIP3 BNIP3L BRP44 BRP44L C18orf55 C22orf29 C22orf32 C2orf18 C4orf49 CAMK2A CASP8 CCDC109B CCDC90A CHCHD4 CNP CPT1A CPT1B CPT2 DNAJC19 DNLZ DYNLL1 DYNLL2 E2F1 EFHA1 EYA2 FBXO7 FIS1 FLVCR1 FXC1 GDAP1 GSK3A GSK3B GZMB HEBP2 HK2 HSP90AA1 HSPA1A HSPA4 IMMP1L IMMP2L KIAA1279 LOC100652748 LOC347411 MAPK8 MAPK9 MCU MFF MFN2 MICU1 MIPEP MOAP1 MPV17L MRPL18 MRS2 MTCH2 MTERFD2 MTX1 MTX2 MTX3 MUL1 NAIF1 NDUFA13 NMT1 NOL3 PAM16 PDE2A PMAIP1 PMPCA PNPT1 PPIF PPP1R13B PPP3CC PPP3R1 PRKAA2 PRKAB2 PRKAG2 PSEN1 RHOT1 RHOT2 SAMM50 SFN SIVA1 SLC1A3 SLC24A6 SLC25A10 SLC25A12 SLC25A13 SLC25A14 SLC25A15 SLC25A20 SLC25A24 SLC25A30 SLC25A32 SLC25A33 SLC25A37 SLC25A5 SLC25A6 SLC8A3 SLC9A1 STARD3 STAT3 STOML2 TFDP1 THEM4 TIMM10 TIMM13 TIMM17A TIMM17B TIMM22 TIMM23 TIMM44 TIMM50 TIMM8B TIMM9 TMEM102 TOMM20 TOMM20L TOMM22 TOMM34 TOMM40 TOMM40L TOMM5 TOMM7 TOMM70A TP53 TP53BP2 TP63 TP73 TRNT1 TSPO TST UCP1 UCP2 UCP3 VDAC1 YWHAB YWHAE YWHAG YWHAH YWHAQ YWHAZ ZNF205 SLC25A1 TAMM41* |
| **Cellular response to Oxidative Stress** |

| |
|---|
| *ABCC2 ABL1 ADA ADAM9 ADIPOQ ADNP2 ADPRHL2 AGER AIF1 AIFM1 AKR1B1 AKR1C3 AKT1 ALAD ALDH3A2 ALDH3B1 ALS2 ANGPTL7 ANKRD2 ANXA1 APEX1 APOA4 APOD APOE APP APTX AQP1 AREG ARG1 ARL6IP5 ARNT ARNTL ATF4 ATOX1 ATP13A2 ATP2A2 ATP7A ATRN BAD BAK1 BCL2 BMP4 BNIP3 C20orf111 CA3 CASP3 CAT CBX8 CCL19 CCL5 CCNA2 CCR7 CCS CD36 CD38 CDK1 CDK2 CHD6 CHRNA4 CHUK CLN8 COL1A1 CPEB2 CRYAB CRYGD CYBA CYBB CYCS CYGB CYP1B1 CYP2E1 DDIT3 DGKK DHCR24 DHRS2 DIABLO DNM2 DPEP1 DUOX1 DUOX2 DUSP1 ECT2 EDN1 EEF2 EGLN1 ENDOG EP300 EPAS1 EPX ERCC1 ERCC2 ERCC3 ERCC6 ERCC8 ERO1L ETFDH ETS1 ETV5 EZH2 F3 FABP1 FAS FER FGF8 FKBP1B FN1 FOS FOSL1 FOXO1 FOXO3 FXN G6PD GAB1 GCLC GCLM GJA3 GJB2 GLRX2 GNAO1 GPX1 GPX2 GPX3 GPX4 GPX5 GPX6 GPX7 GPX8 GSK3B GSR GSS GSTP1 GUCY1B3 HAO1 HBA1 HBA2 HBB HDAC2 HDAC6 HIF1A HMOX1 HNRNPD HP HSPA1A HSPA1B HTRA2 HYAL1 HYAL2 IDH1 IL18BP IL18RAP IL1B IL1R1 IL6 IMPACT IPCEF1 JAK2 JUN KAT2B KCNA5 KCNC2 KDM6B KLF2 KLF4 KLF6 KPNA4 KRT1 LDHA LIAS LONP1 LPO LRRK2 MAP3K5 MAPK7 MAPT MB MBL2 MDM2 MELK MGMT MGST1 MICB MMP14 MMP3 MPO MPV17 MSRA MSRB2 MSRB3 MST4 MT3 MTF1 MTR MUTYH NAPRT1 NDUFA12 NDUFA6 NDUFB4 NDUFS2 NDUFS8 NEFH NEIL1 NET1 NFE2L1 NFE2L2 NFKB1 NGFR NOS1 NOS3 NOX4 NOX5 NQO1 NR4A2 NR4A3 NUDT1 NUDT2 OGG1 OLR1 OXR1 OXSR1 P4HB PARK2 PARK7 PARP1 PAX2 PCGF2 PDCD10 PDGFD PDGFRA PDGFRB PDK1 PDK2 PDLIM1 PENK PINK1 PKD2 PLA2G4A PLA2R1 PLEKHA1 PLK3 PML PNKP PNPT1 PON2 PPARGC1A PPARGC1B PPIF PPP1R15B PPP2CB PPP5C PRDX1 PRDX2 PRDX3 PRDX5 PRDX6 PRKAA1 PRKCD PRKD1 PRKRA PRNP PRODH PSEN1 PSIP1 PSMB5 PTGS1 PTGS2 PTK2B PTPRK PTPRN PXDN PXDNL PXN PYCR1 PYCR2 RAD52 RBM11 RELA RGS14 RHOB ROMO1 RPS3 RRM2B S100A7 SCARA3 SCGB1A1 SDC1 SELK SELS SEPN1 SEPP1 SEPX1 SETX SGK2 SHC1 SIN3A SIRT1 SIRT2 SLC11A2 SLC23A2 SLC25A24 SLC7A11 SLC8A1 SNCA SOD1 SOD2 SOD3 SPHK1 SRC SRXN1 STAR STAT6 STAU1 STC2 STK24 STK25 STX2 STX4 TACR1 TAT TMEM161A TNFAIP3 TOR1A TP53 TP53INP1 TPM1 TPO TRAF2 TRPA1 TRPC6 TRPM2 TXN TXN2 TXNDC2 TXNDC3 TXNDC8 TXNIP TXNL1 TXNRD1 TXNRD2 UCN UCP2 UCP3 VKORC1L1 VNN1 VRK2 WNT16 WRN XBP1 XPA ZC3H12A ZNF277 ZNF580 ZNF622* |
| **Apoptotic Mitochondrial Changes** |
| *BAK1 BAX BBC3 BCL2L1 BID CDKN2A DNM1L GPX1 IFI6 PMAIP1 SFN ACAA2 AIFM2 AKT1 ATF2 ATG3 ATP2A1 BAD BCL2 BCL2A1 BCL2L11 BIK BLOC1S2 BMF BNIP3 BNIP3L C22orf29 CAMK2A CCK CLU ERBB4 EYA2 FIS1 GCLC GCLM GGCT HK2 IFIT2 JTB JUN KIAA1967 MAPK9 MCL1 MFF MOAP1 MTCH2 NAIF1 NDUFS1 NOL3 PPIF PPP2CB RHOT1 RHOT2 SLC25A4 SOD2 ST20 THEM4 TIMM50* |
| **Calcium Homeostasis and Transport** |
| *ANXA6 ATP2A1 BCAP31 C22orf32 CCDC109B CCDC90A DISC1 EFHA1 FIS1 IMMT MCU MICU1 RAP1GDS1 SLC24A6 SLC8A3 TGM2 BHLHA15 STOML2 VDAC1* |
| **Mitochondrial Fission and regulation** |
| *COX10 DNM1L FAM54A FAM54B FIS1 GDAP1 GGNBP1 MFF MTFP1 MTFR1 MUL1 OPA1 PARK2 SLC25A46 SMCR7L BNIP3 DCN DDHD1 DDHD2 KDR MARCH5 PINK1 SMCR7 DRP1* |
| **Mitochondrial Fusion** |
| *AFG3L2 BAK1 BAX BCL2A1 CHCHD3 FAM73A FAM73B FIS1 GDAP1 MFF MFN1 MFN2 OPA1 PLD6 SMCR7 SMCR7L ST20 STOML2 USP30* |
| **Mitophagy and its regulation** |

| |
|---|
| *ACIN1 ACTRT1 ADAMTS7 AKR1E2 ALPK1 AMBRA1 ANXA5 ASB2 ATG13 ATG14 ATP1B1 ATPAF1-AS1 ATPIF1 BECN1 BLOC1S1 BMP2KL BOC C11orf41 C5 C8orf59 CA7 CALCB CAPS CD163L1 CD93 CDC37 CHAF1B CHST3 CLVS1 COX8A CRNKL1 CSPG5 DKKL1 DNAAF2 DPF3 DZANK1 EIF2S1 FABP1 FAM131B FAM13B FAM176B FANCC FANCF FCGR3B FGFBP1 GABRA5 GDF5 GMIP HAPLN1 HCAR1 HDAC6 HK2 HPR HSF2BP IPPK IST1 ITPKC KCNK3 KRCC1 KRT15 KRT73 LARP1B LENG9 LMCD1 LSM4 MAP1A MAP2K1 MAP3K12 MBD5 MDH1 MEX3C MRPS10 MRPS2 MSTN MYH11 MYLK MYOM1 NDUFB9 NEFM NME2 NR2C2 NTHL1 NUP93 OBSCN P2RX5 PARK2 PDK1 PEX13 PEX3 PFKP PGK2 PHYHIP PI4K2A PIK3CA PINK1 PLOD2 PNPO PPY PRKD2 PRKG1 REP15 RFWD3 RIMS3 SERPINB10 SFRP4 SLC1A3 SLC1A4 SLC22A3 SLC35B3 SLC35C1 SLC37A4 SLC6A1 SLCO1A2 SMURF1 SNRPB SNRPD1 SNRPF SNTG1 SQSTM1 STAT2 STK32A STOM SUPT3H TBC1D5 TMEM203 TMEM39A TMEM39B TOMM7 TREM1 TXLNA YIPF1 ZCCHC17 ZNF189 ZNF593 ACTL6A ADRB2 ATG5 ATG7 ATP13A2 BNIP3L C12orf5 CSNK2A2 CTSK CTTN DNM1L FBXW7 FZD5 GSK3A HAX1 HIF1A HTRA2 KAT2A MAP1LC3B PARK7 PARL POLR3A RPL28 RUVBL1 SLC17A9 SREBF1 SREBF2 TSPO U2AF1 U2AF2 USP36 VDAC1 VPS13C WBP11 WDR75 ZBTB17 ZDHHC8* |
| **Mitonuclear Crosstalk** |
| *CAMK4 CAMKK2 CREB ESRRA ESRRB ESRRG GABPA MEF2A NCOR1 NRF1 NRIP1 PARP1 PARP2 PPARA PPARG PPARGC1A PPARGC1B PPRC1 PRKAA1 PRKACA PRKACB SIRT1 TP53 ATF2 CEBPG DDIT3 EGR1 KEAP1 MAPK1 MAPK8 MAPK9 MAPK10 MTOR NFATC1 NFE2L2 NFKB1 NFKB2 PPP3CC* |
| **Fatty Acid Beta oxidation** |
| *ACADL ACADM ACADS ACADVL DECR1 ECHS1 ECI1 HADH HADHA HADHB MCEE MUT PCCA PCCB* |
| **Mitochondrial Membrane Potential regulation** |
| *ABL1 ALOX12 ARL6IP5 ATPIF1 BAD BAK1 BAX BCL2 BCL2L1 BCO2 C22orf29 CASP1 CCK CDKN2A CLIC1 DCN GCLC GCLM GNB2L1 GSK3B HEBP2 HSH2D IFI6 KDR LRRK2 MLLT11 MUL1 MYOC NDUFS1 NGFR OPRD1 P2RX7 PANK2 PARK2 PARK7 PID1 PINK1 PMAIP1 PPP2R3C PRDX3 PRELID1 PYCR1 SLC25A33 SLC25A36 SOD1 SPG20 SRC STOML2 STOX1 TSPO TUSC2 UBB UCP2 VCP* |

#### F. Phenotypic variance explained by each mitochondrial pathway

The amount of phenotypic variance explained by the genetic contribution of each mitochondrial pathway (represented by the pseudo-$r^2$ metric proposed by [3]) was calculated using PRSet. Age, sex, and the first three principal components were included as covariates. This provided a sample-wide summary of the predictive power of each pathway-PRS [4].

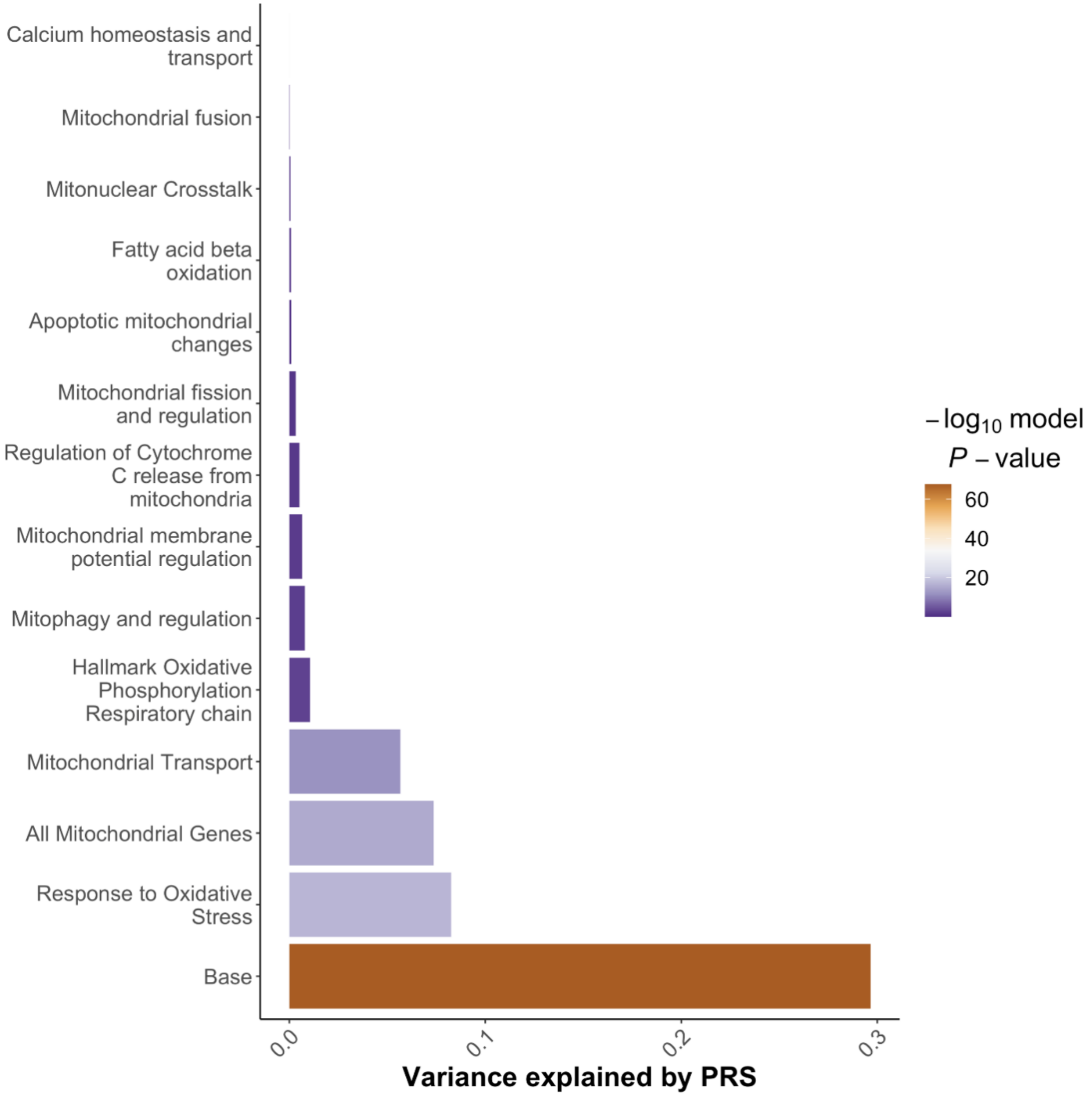

***Fig. F.1 Variance in Alzheimer's phenotype explained by the polygenic risk score for pathways and gene regions****. The colour gradient represents the significance level of competitive empirical p-value for AD case – CN control regression.*

The amount of phenotypic variance explained by the genetic contribution of each mitochondrial pathway (represented by the pseudo-$r^2$ metric proposed by [3]) was calculated adding age, sex, and first three principal components as covariates in the model. The highest amount of AD phenotypic variance in the model is explained by the whole nuclear genome PRS (~30%) followed by OXTRESS pathway (~8%). The variance explained by complete nMT-DNA, mitochondrial transport, OXPHOS, mitophagy and regulation, and mtΔΨ regulation pathway-PRS was 7%, 5%, 1.5%, 1% and 0.09% respectively.
